# Modeling centrosome positioning by actin–microtubule crosstalk and actin anchoring at the cell membrane

**DOI:** 10.64898/2026.10.05.756642

**Authors:** Prasanta Kumar Biswas, Utsa Dey, Apurba Sarkar, Raja Paul

## Abstract

Centrosome positioning is governed by mechanical interactions between the actin and microtubule cytoskeletons, yet the contribution of different actin architectures and their attachment to the cell membrane remains unclear. We use numerical simulations to study a centrosomal array of microtubules interacting with cortical, bulk, and centrosome-associated actin networks, including passive cross-linkers, motor proteins, and ERM-mediated coupling of actin to the cell membrane. We find that bulk and centrosome-associated actin alone are insufficient for robust centrosome centering, whereas sufficiently dense cortical actin promotes centering through steric interactions with growing microtubules. ERM-mediated attachment of cortical actin to the cell membrane enhances this effect, allowing stable centering at lower actin densities. In a two-centrosome system, microtubule interactions generate a spindle-like structure, while the organization of actin and its membrane attachment through ERM regulate centrosome separation. Antagonistic microtubule motors further limit centrosome separation and establish a finite spindle length. These results demonstrate that ERM-mediated coupling of actin to the cell membrane provides an important mechanical link between the actin cortex and microtubule-driven centrosome positioning, influencing both centrosome centering and spindle geometry.

## I. INTRODUCTION

The centrosome (CS) is the primary microtubule-organizing center in animal cells and plays an important role in organizing the intracellular space [1]. Its position is involved in several cellular processes, including cell polarization, migration, immune synapse formation, intracellular trafficking, and mitotic spindle assembly [2– 5]. Proper positioning of the CS is therefore essential for normal cellular function, while defects in CS dynamics have been linked to developmental abnormalities and disease [6, 7].

Microtubules provide a major source of mechanical forces that position the CS. Polymerization of microtubules can generate pushing forces that displace the CS from the cell cortex [8]. Conversely, cortical motors such as dynein can pull on astral microtubules near the cell boundary and thereby exert forces that position the CS toward the cortex [9, 10]. The balance between these opposing forces can establish a stable CS position within the cell [11]. Experimental and theoretical studies have provided substantial insight into these microtubule-based mechanisms of CS positioning [12].

Actin provides another mechanical network that can contribute to CS positioning [13]. Cortical actin has been implicated in the mechanical regulation of nuclear and CS movements, and actin structures associated with the CS have also been observed [14]. Importantly, actin is organized into distinct architectures within the cell, including bulk cytoplasmic networks, cortical networks [15], and CS-associated filaments [16]. These networks differ in their spatial distribution and in their mechanical interactions with microtubules and the cell boundary. Their contributions to CS positioning may therefore depend strongly on their organization.

This dependence on actin architecture was recently demonstrated in a cell-free reconstitution study in which a microtubule aster interacted with differently organized actin networks confined within cell-sized microwells [17]. Simulations incorporating steric interactions between the two cytoskeletal networks showed that bulk actin restricted microtubule displacement but did not produce CS centering, whereas dense cortical actin near the cell boundary promoted centering through a balanced distribution of pushing forces. However, the actin filaments in these simulations remained fixed at their initial nucleation sites. Such an assumption does not account for the remodeling and displacement of actin networks in cells, where membrane attachment and actin-binding proteins can stabilize the network [18]. Thus, the mechanical coupling between actin networks and the cell cortex provides an important factor to consider when examining actin-dependent CS positioning.

The Ezrin-Radixin-Moesin (ERM) proteins are key membrane-actin linkers that can provide such coupling by connecting filamentous actin to the plasma membrane [19, 20]. ERM-mediated attachment can stabilize cortical actin and facilitate force transmission between the actin network and the cell cortex, while also contributing to cell shape, mechanics, and signaling [21]. Despite this established mechanical role, how ERM-mediated actin anchoring affects CS positioning has not been systematically investigated.

The mechanical state of the actin network is also affected by passive cross-linking proteins and active myosin motors. Passive cross-linkers connect neighboring actin filaments, whereas myosin motors generate contractile forces and drive network rearrangement [22, 23]. Whether these changes in actin connectivity and contractility are important for CS positioning, or whether cortical anchoring provides the dominant mechanical contribution, remains unclear.

The role of actin organization becomes even more relevant when considering two CSs, as encountered during mitotic spindle assembly. The two CSs form the microtubule-organizing centers of the bipolar spindle, and their separation must be coordinated to maintain spindle architecture and function [18, 24]. While microtubule interactions are known to contribute to spindle organization, the extent to which different actin architectures and their coupling to the cell cortex influence CS separation remains unresolved.

A wide range of factors has been considered previously to study centrosome positioning [25], including MT pushing forces against the cell and nuclear boundaries [9, 26], pushing forces arising from MT-organelle interactions [11], dynein-mediated pulling from the cortex and nuclear surface [9, 11, 27, 28], cytoplasmic dynein-mediated pulling along the MT lattice [11, 29], and actomyosin-driven centripetal flows [30]. Recent experiments have also highlighted a role for MTs and cytoplasmic dynein-dependent cell-shape remodeling around the centrosome in promoting its central positioning [11, 30]. Although each of these mechanisms can contribute to centrosome positioning, how the spatiotemporal interplay between actin and microtubule networks, particularly in the presence of motor proteins and cross-linkers, regulates centrosome organization remains poorly understood. For example, microtubules can generate pushing forces against the cell membrane only if their tips experience sufficient friction as they slide along it. A dense, mechanically coupled actin network can stabilize an off-center centrosome by resisting microtubule-generated forces, whereas a sparse or weakly anchored network may allow continued centrosome displacement [30]. Can motor proteins within the actin network or passive cross-linkers compensate for weak cortical anchoring and restore centrosome centering? More broadly, how do these actin-generated mechanical contributions interact with microtubule forces to maintain the positioning of a centrosome near the cell center during interphase and to organize duplicated centrosomes into a bipolar, spindle-like structure during mammalian mitosis?

In this work, we use *Cytosim*-based [31] simulations to investigate the mechanical role of bulk, cortical, and centrosomal actin networks in CS positioning. We examine how ERM-mediated cortical anchoring and ERM turnover affect CS centering and further determine the contributions of passive actin cross-linkers and myosin motors. We then extend the model to two CSs to examine how actin organization, cortical anchoring, and microtubule interactions regulate CS separation and spindle structure.

We find that CS positioning depends strongly on actin organization and its coupling to the cortex. Bulk and centrosomal actin require ERM-mediated anchoring for efficient centering, whereas sufficiently dense cortical actin can center the CS without ERM. ERM anchoring further enhances cortical-actin-dependent centering, while actin cross-linkers and myosin motors have limited effects. In the two-CS system, microtubule steric interactions generate a spindle-like structure, while actin density and ERM anchoring regulate CS-CS separation. Antagonistic microtubule motors establish a finite bipolar spindle and, together with ERM-anchored actin, further control spindle length. Thus, cortical anchoring provides a key mechanical link through which actin organization regulates both CS positioning and spindle geometry.

### II. MODEL AND SIMULATION

We consider a minimal two-dimensional model containing microtubules nucleated from centrosomes (CSs), actin filaments, ERM linkers, molecular motors, and actin cross-linkers, all confined within a circular cell boundary as shown in Fig. 1. Simulations were performed using the open-source cytoskeleton simulation package *Cytosim* [31].

**FIG 1:**
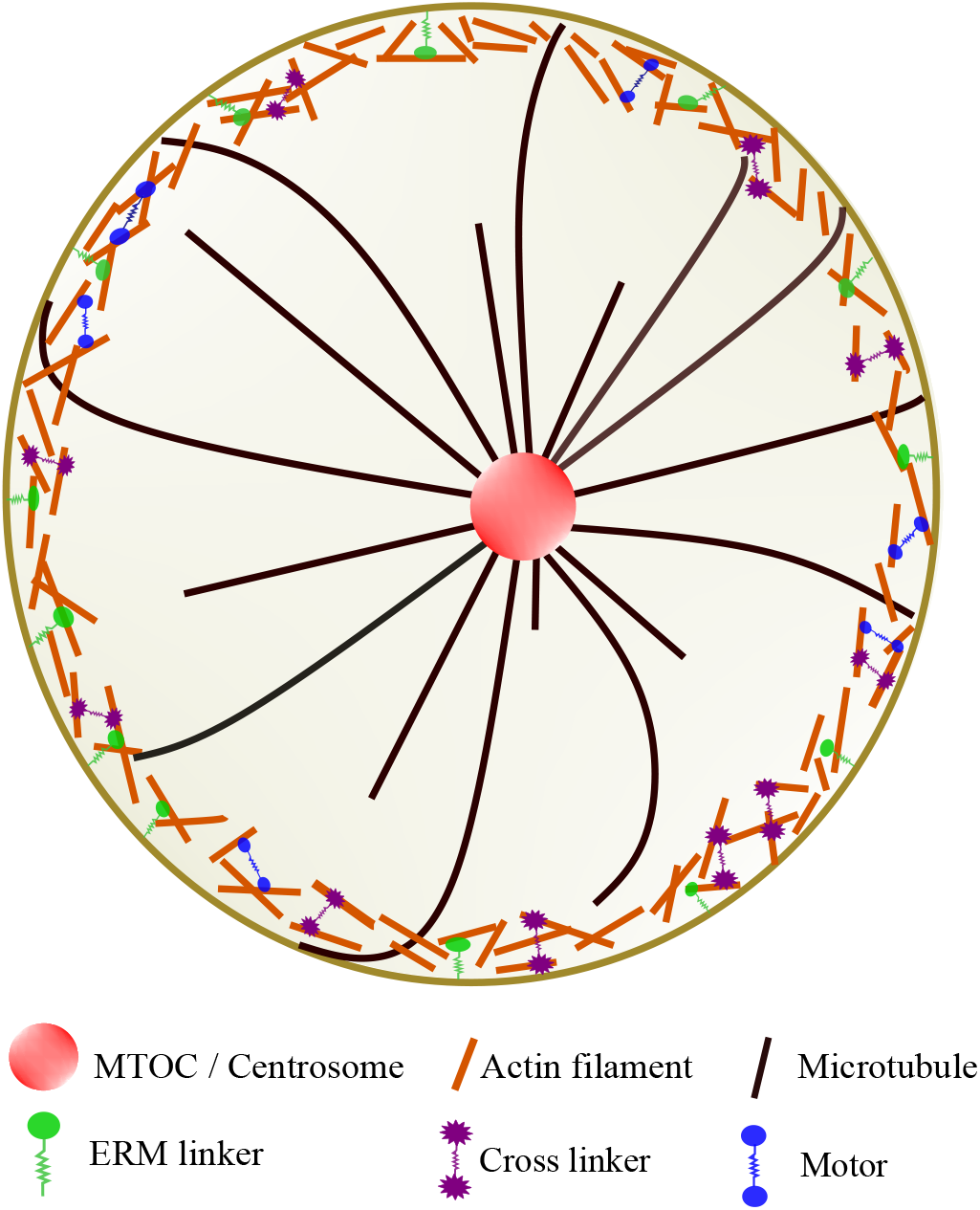
Schematic illustration of the model cell highlighting the organization of centrosomal microtubules, cortical actins, ERM linkers, molecular motors and cross-linkers.

Microtubules and actin filaments were modeled as semi-flexible polymers. Microtubules were nucleated from the CSs with their minus ends fixed at the CS, while their plus ends underwent dynamic instability [32]. Transitions between growth and shrinkage occurred stochastically with catastrophe and rescue frequencies, respectively. Microtubules changed length during these phases with the corresponding growth and shrinkage velocities, while actin filaments were modeled with fixed lengths.

Each CS was represented by a rigid bead with an effective viscosity chosen to reproduce the experimentally measured CS mobility [12].

ERM linkers were modeled as passive membrane-actin linkers. One end of each linker was permanently attached to the cell boundary, while the other end could bind to and detach from nearby actin filaments within a specified binding range. Upon binding, the linker remained immobile and acted as an elastic spring, thereby coupling the actin network mechanically to the cell boundary and transmitting forces between them.

Cytoplasmic active and passive cross-linkers were represented as two-headed elements connected by an elastic spring. Each head could stochastically bind to and detach from actin filaments. Once bound, the heads of active cross-linkers moved toward a prescribed filament end, whereas passive cross-linkers remained stationary. Both types of cross-linkers generated elastic forces that brought the associated actin filaments closer together.

Microtubule-actin interactions were modeled through steric repulsion. Microtubules exerted repulsive forces on actin filaments along their length, while actin filaments exerted repulsive forces on growing microtubule plus ends. These interactions opposed microtubule growth and contributed to the mechanical positioning of the CSs.

The dynamics of all model components were simulated within the *Cytosim* framework using Langevin dynamics. The thermal energy was set to *k*_*B*_*T* = 4.2 pN · nm, and the effective cytoplasmic viscosity was set to 1 pN·s*/µ*m^2^ [33]. Cell confinement was imposed by applying a Hookean restoring force to intracellular components that crossed the prescribed cell boundary. The complete list of model parameters are given in the Appendix A.

## III. RESULTS

### A. Cortical actin networks regulate CS positioning

Cortical actin networks can strongly influence CS positioning. These filaments form a mesh-like layer beneath the plasma membrane and interact mechanically with microtubules, potentially contributing to CS stabilization. To examine this role, we considered four distinct configurations and determined their effects on the steady-state CS position. The results for each case are presented in the following sections.

#### 1. High-density cortical actin promotes CS centering

In our simulations, we systematically varied the number of cortical actin filaments localized near the cell periphery between 500 and 3500 (500, 2000, and 3500) to investigate their effect on CS positioning. Only steric interactions between microtubules and actin filaments were considered, and no membrane-actin cross-linkers were included. Under these conditions, CS positioning was determined solely by passive mechanical interactions among the cytoskeletal components, as illustrated in Figs. 2(a) 2(i).

**FIG 2:**
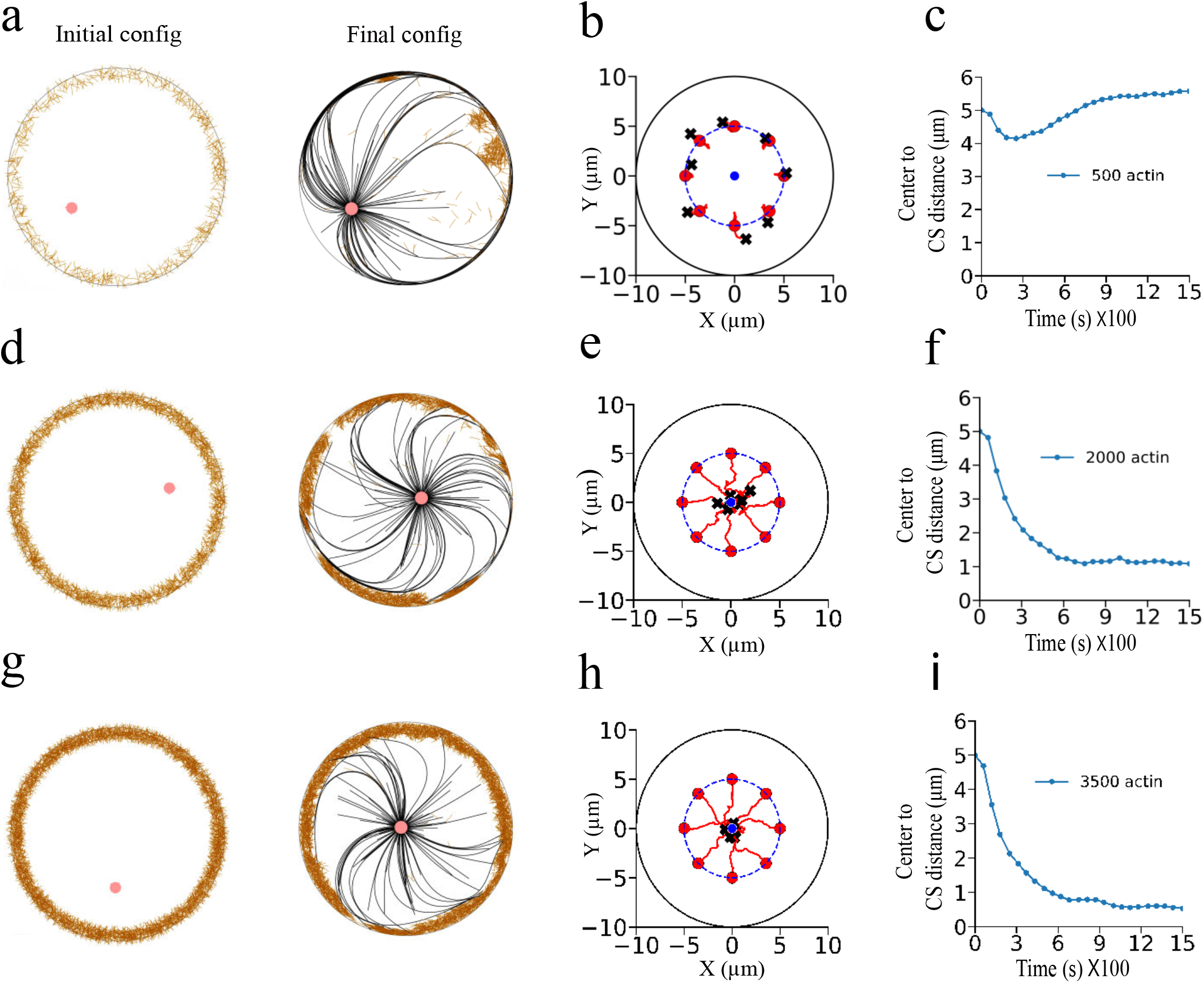
CS positioning with cortical actin and no ERM linkers. (a) Initial and final configurations for 500 cortical actin filaments. (b) CS trajectories for different initial positions along a virtual circle of radius 5 *µ*m; red filled circles and black crosses denote the initial and final CS positions, respectively. (c) Average CS distance from the cell center. At low actin density, growing microtubules sweep actin filaments away from their path, leading to their accumulation in one region and an asymmetric force that displaces the CS from the cell center. (d–e) Initial and final configurations and CS trajectories for 2000 cortical actin filaments. (f) Average CS distance from the cell center. At higher actin density, the actin network restricts microtubule movement along the cell boundary, producing a more balanced force and a CS position closer to the cell center. (g–h) Initial and final configurations and CS trajectories for 3500 cortical actin filaments. (i) Average CS distance from the cell center, showing enhanced CS centering with increasing actin density.

When the cortical actin density was low (~500 filaments), as shown in Fig. 2(a), growing microtubules readily displaced nearby actin filaments through steric interactions and extended to the cell boundary with little resistance. This allowed the microtubules to glide freely along the cell cortex, producing an uneven distribution of pushing forces on the CS. As a result, the restoring force toward the cell center remained weak, and stable CS centering was not achieved. This behavior is reflected in Fig. 2(b), where CSs starting from different initial positions fail to return to the cell center, and in Fig. 2(c), where the CS-cell-center distance remains large throughout the simulation.

Increasing the cortical actin density to 2000 filaments significantly altered the mechanical response of the network. At this density, growing microtubules were no longer able to completely displace cortical actin filaments from their paths and therefore experienced much stronger steric resistance. The reduced microtubule gliding produced a more uniform distribution of pushing forces around the CS, leading to nearly balanced forces from all directions and driving the CS toward the cell center. A representative configuration is shown in Fig. 2(d), where the cortical actin network is locally reorganized through its interactions with microtubules. Correspondingly, CSs released from different initial positions converge toward the cell center, as shown in Figs. 2(e) and 2(f). Increasing the number of cortical actin filaments further to 3500 strengthened this effect, allowing the CS to reach the cell center more rapidly while largely preserving the integrity of the cortical actin network (Figs. 2(g) 2(i))

These results indicate that robust CS centering requires the cortical actin density to exceed a threshold. In our simulations, stable centering first emerged at approximately 2000 cortical actin filaments, whereas lower densities produced only weak or transient centering. Thus, steric interactions between microtubules and cortical actin filaments can generate the restoring forces required for CS centering, provided that the cortical network is sufficiently dense to offer collective mechanical resistance. In the absence of membrane-actin cross-linkers, a dense cortical actin network is therefore necessary to maintain stable CS centering.

#### 2. ERM membrane cross-linkers reduce the cortical actin density required for CS centering

We next introduced ERM proteins into the model and systematically varied the number of cortical actin filaments to 500, 2500, and 3500, as illustrated in Fig. 3(a). ERM proteins (Ezrin-Radixin-Moesin family) are membrane-associated cross-linkers that localize at the inner surface of the plasma membrane and mechanically couple actin filaments to the cell cortex [19, 20]. In contrast to the case containing only cortical actin filaments, where steric interactions between microtubules and actin filaments constituted the dominant mechanical interaction, the presence of ERM linkers enabled robust CS centering at a much lower actin density. In our simulations, approximately 500 cortical actin filaments were sufficient to achieve stable CS centering, as shown in Figs. 3(b) and 3(c). By anchoring cortical actin filaments to the membrane, they suppress the displacement of actin filaments by growing microtubules and thereby stabilize the cortical actin network. This stabilized network provides a mechanically resistant boundary against microtubule-generated forces.

**FIG 3:**
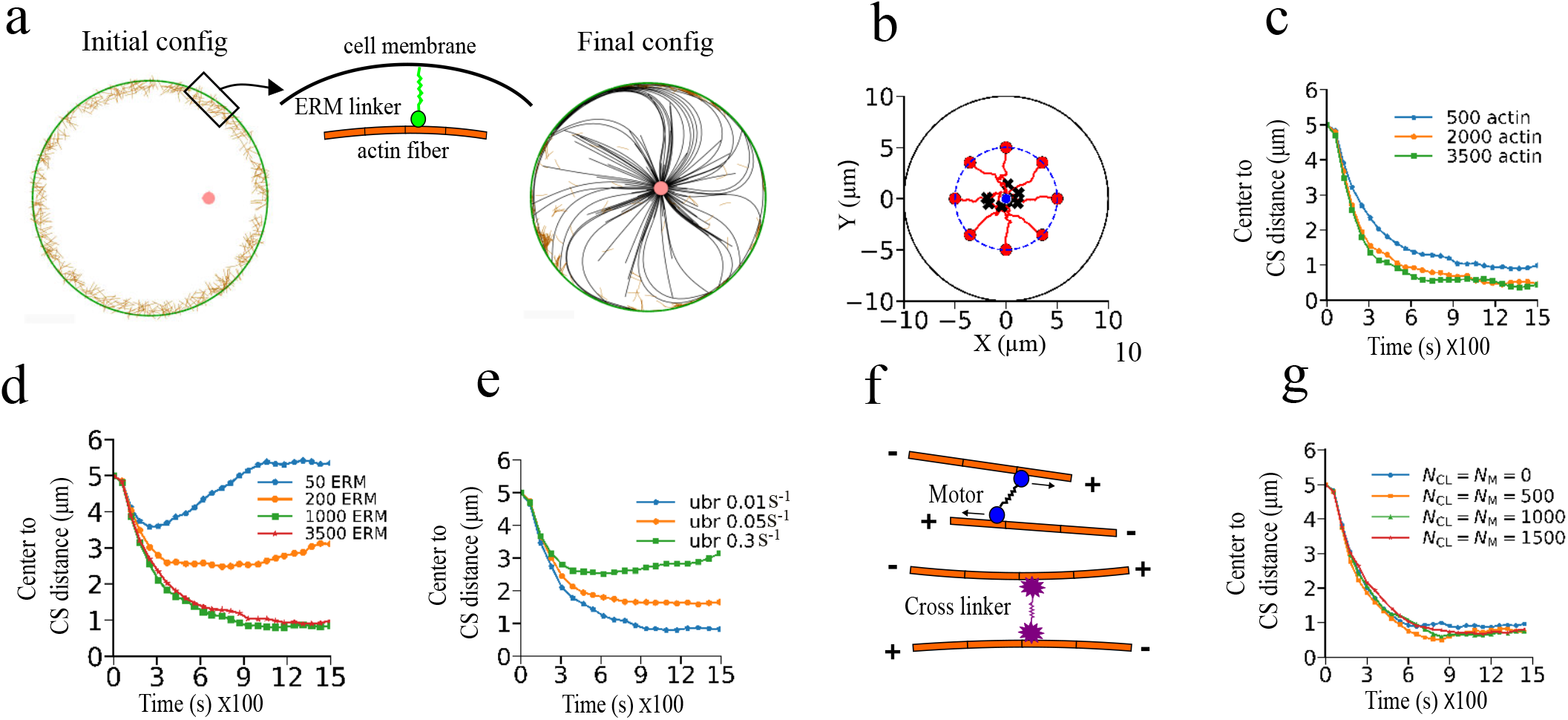
CS positioning with cortical actin and ERM linkers. (a) Initial and final configurations for 500 cortical actin filaments, with a magnified view of ERM-mediated actin–membrane anchoring. (b) CS trajectories from different initial positions along a virtual circle of radius 5 *µ*m; red filled circles and black crosses denote the initial and final CS positions, respectively. (c) Mean CS distance from the cell center for different actin densities, showing robust centering at 500 filaments. (d) CS centering versus ERM linker number at a fixed unbinding rate of 0.01 *s*^−1^. (e) CS centering versus ERM linker unbinding rate for 1000 linkers, showing that efficient centering requires sufficient linker density and slow turnover. (f) Schematic of passive actin cross-linkers and active myosin motors. (g) Mean CS distance for different numbers of cross-linkers (*N*_*CL*_) and myosin motors (*N*_*M*_), showing limited effects on CS positioning. Panels (d), (e), and (g) use 500 cortical actin filaments.

This effect is evident from the CS dynamics. In the absence of ERM linkers, a cortical actin number of ~500 filaments was insufficient for stable CS centering because growing microtubules readily displaced the loosely connected cortical network. Consequently, microtubules glided along the cell boundary, producing an asymmetric distribution of pushing forces around the CS. When ERM linkers were introduced, the same actin density became sufficient for robust CS centering because the anchored cortical network resisted microtubule-induced rearrangements and restricted microtubule gliding. As a result, the pushing forces became nearly symmetric, driving the CS toward the cell center.

When the cortical actin number was reduced further to ~200 filaments or less, CS centering was again lost, even in the presence of ERM linkers. Although the actin filaments remained anchored to the membrane, the network was too sparse to provide sufficient mechanical resistance against the growing microtubules (Appendix B). Increasing the cortical actin count beyond ~500 progressively strengthened the anchored network, suppressing microtubule gliding and producing a more uniform distribution of forces on the CS. These results show that efficient CS centering requires both ERM-mediated cortical anchoring and a sufficiently dense cortical actin network. ERM anchoring stabilizes the cortical layer, while the actin density determines the magnitude of the restoring forces generated through steric interactions with microtubules.

#### 3. ERM linker properties influence CS positioning

With the inclusion of ERM protein linkers, it is natural to ask how their number and binding properties influence CS centering. In the previous section, we found that, with 3500 ERM linkers, a minimum of 500 cortical actin filaments was sufficient to achieve stable CS centering. We therefore fixed the number of cortical actin filaments at 500 and systematically varied the ERM linker parameters to examine their effect on CS positioning. We first investigated the influence of ERM linker density by varying the number of ERM linkers while keeping their unbinding rate fixed. The unbinding rate characterizes the rate at which an ERM linker detaches from an actin filament.

Simulations were performed with 50, 200, 1000, and 3500 ERM linkers while maintaining the unbinding rate at 0.01 *s*^−1^, as shown in Fig. 3(d). With only 50 ERM linkers, the CS failed to move to the cell center when released from an off-center position. At this low linker density, only a small fraction of the cortical actin network was anchored to the cell membrane, resulting in weak mechanical coupling between the membrane and the cortical actin network. Consequently, the cortical actin layer could not provide sufficient resistance to microtubules gliding along the cell cortex, leading to asymmetric force transmission and CS decentering. Increasing the number of ERM linkers to 200 produced partial CS centering, indicating improved cortical anchoring and more effective transmission of microtubule-generated forces. Stable CS centering was achieved with 1000 ERM linkers, suggesting that this number was sufficient to anchor the cortical actin network over most of the cell boundary. Further increasing the number of ERM linkers to 3500 produced essentially the same behavior, indicating that approximately 1000 ERM linkers were sufficient for robust CS centering.

Next, we investigated the effect of the ERM linker unbinding rate while keeping the number of ERM linkers fixed at 1000. Simulations were performed for unbinding rates of 0.01 *s*^−1^, 0.05 *s*^−1^, and 0.3 *s*^−1^, as shown in Fig. 3(e). We found that CS centering progressively deteriorated as the ERM unbinding rate increased. At an unbinding rate of 0.01 *s*^−1^, ERM linkers remained attached to actin filaments for long periods, producing a strong and stable mechanical connection between the cortical actin network and the cell boundary. This persistent anchoring enabled efficient transmission of restoring forces and promoted robust CS centering. At an intermediate unbinding rate of 0.05 *s*^−1^, more frequent detachment events weakened the cortical anchoring, resulting in less stable CS positioning. When the unbinding rate was increased to 0.3 *s*^−1^, ERM linkers detached rapidly from actin filaments and failed to maintain a stable connection between the cortical actin network and the cell boundary. Consequently, microtubules were again able to glide along the cell cortex with reduced resistance, leading to asymmetric force transmission and gradual CS decentering.

#### 4. CS positioning in the presence of active and passive actin cross-linkers

The stable CS centering discussed in the previous sections relies on the mechanical integrity of the cortical actin network upon interaction with the cell membrane. We next investigated whether modifying the internal connectivity of this network through active and passive actin cross-linkers influences CS positioning. Passive actin cross-linkers (e.g., *α* actinin, filamin, fascin) mechanically connect neighboring actin filaments and primarily reinforce the structural integrity of the actin network without generating active forces [34, 35]. In contrast, active actin cross-linkers, such as myosin motors, not only bind actin filaments but also generate ATP-driven contractile forces that continuously remodel the actin network and produce internal stresses [36, 37] (see Fig. 3(f)).

To examine the influence of these cross-linkers, simulations were performed in the presence of a fixed number of 3500 ERM linkers. As shown in Fig. 3(g), introducing passive and active cross-linkers produced little change in CS positioning. Once the cortical actin network is firmly anchored to the cell boundary through ERM-mediated membrane-actin coupling, efficient CS centering is already established. The ERM linkers provide stable mechanical coupling between the cortical actin network and the membrane, allowing microtubule-generated forces to be transmitted effectively to the cell cortex. Under these conditions, the additional network connectivity provided by passive cross-linkers and the contractile activity generated by myosin motors do not significantly modify the global force balance acting on the CS. Instead, these components mainly reorganize the local actin network without substantially affecting the restoring forces responsible for CS centering. Consequently, robust CS centering persists over the range of passive cross-linker and myosin motor numbers considered.

We next examined the same system in the absence of ERM linkers. As discussed previously, a dense cortical actin network consisting of ~2000 filaments was sufficient to achieve stable CS centering through steric interactions alone. However, the addition of active and passive actin cross-linkers under these conditions resulted in CS decentering. Without ERM-mediated anchoring, the cortical actin network is free to reorganize in response to microtubule-generated forces. The increased network connectivity introduced by passive cross-linkers and the contractile activity of myosin motors promote local actin rearrangements and generate asymmetric internal stresses, which disrupt the symmetric transmission of forces to the CS. As a result, the restoring forces required for stable CS centering are weakened, leading to CS decentering (Appendix. C).

### B. Influence of bulk actin networks on CS positioning

In this section, we investigate the role of bulk actin in CS positioning using a systematic approach similar to that adopted for cortical actin networks. Bulk actin consists of actin filaments distributed throughout the cytoplasm, where they form an internal cytoskeletal network that contributes to intracellular force transmission and organization. Through its mechanical interactions with microtubules, this network can also regulate CS positioning.

We first examine CS positioning in the presence of bulk actin alone and then introduce ERM linkers to investigate the effect of membrane-actin coupling. We subsequently vary the ERM linker density and unbinding rate, followed by the inclusion of active myosin motors and passive actin cross-linkers to evaluate the effects of network connectivity and contractility. This systematic analysis enables us to assess the contribution of each component to CS positioning.

#### 1. Bulk actin alone fails to achieve CS centering

CS centering was not observed for any of the bulk actin densities investigated (500, 2000 and 3500 total actin filaments) in the absence of cross-linkers and motor proteins. Instead, the CS consistently remained in an off-center position (see Figs. 4(a)-4(c)).

**FIG 4:**
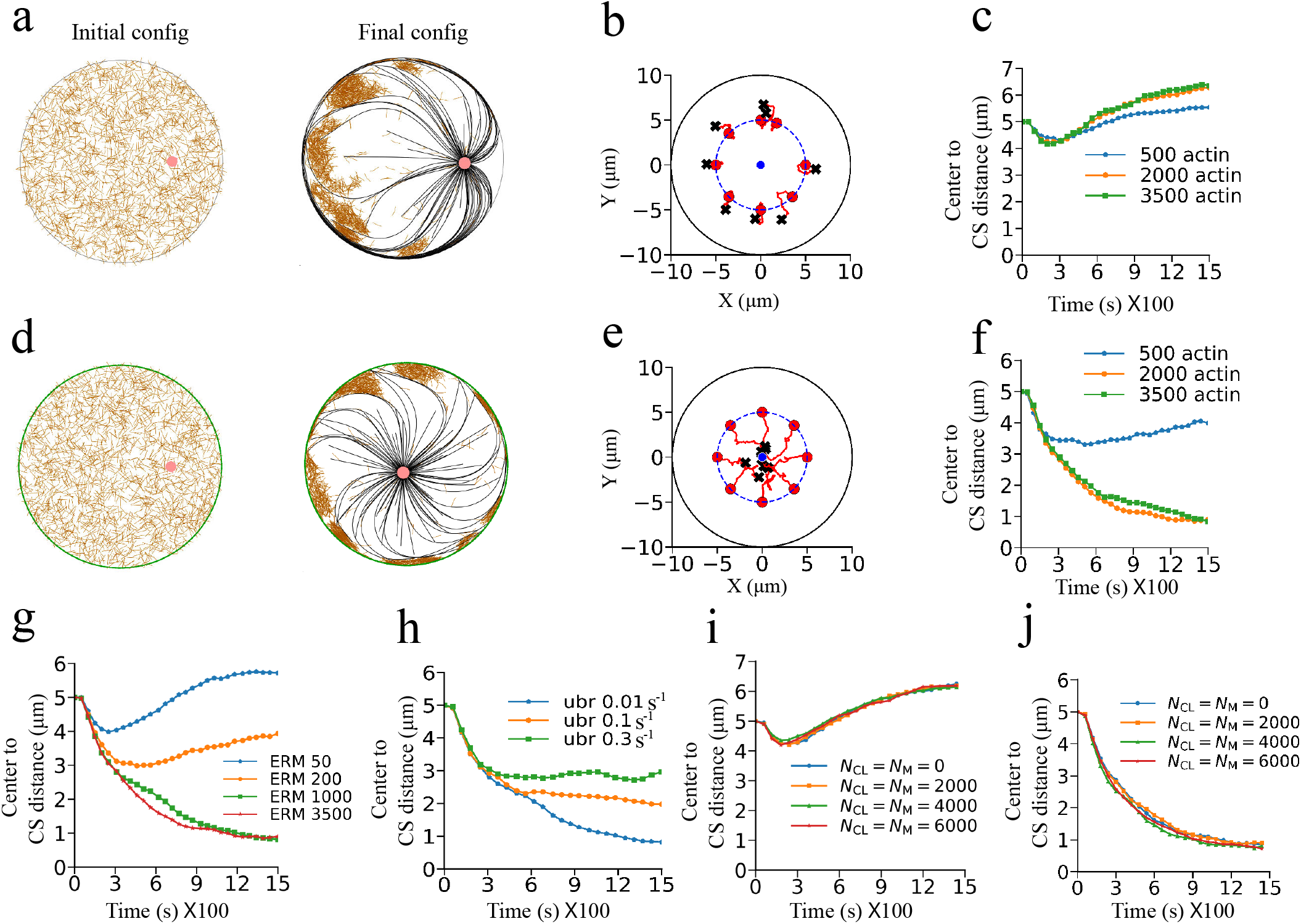
CS positioning in bulk actin networks. (a–c) Without ERM linkers: (a) initial and final configurations for 2000 bulk actin filaments, (b) CS trajectories from initial positions on a virtual circle of radius 5 *µ*m. (c) Mean CS distance from the cell center for different actin densities centering is not possible at any actin density. (d–f) Corresponding results with ERM linkers. (f) Showing at higher bulk actin density ERM promotes CS centering. (g) Mean CS distance for different ERM linker numbers at an unbinding rate of 0.01 *s*^−1^ showing a sufficient number of ERM is needed to get stable centering of CS, and (h) for different unbinding rates with 1000 ERM linkers demonstrating at higher unbinding rate CS fails to attain a stable position near cell center. (i,j) Mean CS distance for varying numbers of passive cross-linkers and myosin motors without and with ERM linkers, respectively, for 2000 bulk actin filaments, demonstrating the importance of ERM-mediated anchoring for CS centering.

As microtubules polymerized and extended from the CS, they displaced bulk actin filaments through steric interactions. Microtubules that reached the cell boundary first progressively cleared actin filaments from their growth paths, while those arriving later displaced the actin network around the cell periphery. This process produced an asymmetric redistribution of bulk actin, leading to its accumulation on one side of the cell (see Figs. 4(a)).

The resulting asymmetry in the actin network generated an uneven mechanical environment around the CS. Microtubules growing through actin-rich regions experienced stronger steric resistance than those on the opposite side, giving rise to an imbalance in the forces acting on the CS. Consequently, the CS was displaced away from the actin-rich region and settled into an off-center position. This behavior persisted across all bulk actin densities examined, indicating that bulk actin alone, irrespective of its density, is insufficient to produce stable CS centering.

### 2. ERM-mediated bulk actin anchoring rescues CS centering

The behavior of the bulk actin network changed substantially upon introducing ERM linkers (~3500) along the cell cortex. As microtubules polymerized, they continued to displace bulk actin filaments through steric interactions. However, once actin filaments reached the cell boundary, they became anchored to the cortex by ERM linkers. This prevented the asymmetric redistribution of bulk actin observed in the absence of ERM-mediated anchoring and established a more uniform actin organization around the cell periphery.

The anchored actin network provided balanced steric resistance to growing microtubules, resulting in a more symmetric distribution of forces on the CS. Consequently, robust CS centering was achieved for 2000 and 3500 bulk actin filaments, as shown in Figs. 4(d)-4(f). In contrast, CS centering was not observed for 500 bulk actin filaments, even in the presence of ERM linkers. At this low density, the actin network was too sparse to provide sufficient cortical coverage for balanced force transmission and failed to establish stable CS centering.

#### 3. ERM linker density and turnover regulate CS positioning

We next investigated how the density and turnover of ERM linkers influence CS positioning in bulk actin networks.

We first varied the number of ERM linkers (50, 200, 1000, and 3500) while keeping the unbinding rate fixed at 0.01 *s*^−1^, as shown in Fig. 4(g). With a low number of ERM linkers (50 and 200), CS centering was not achieved because cortical coverage was insufficient to anchor the bulk actin network uniformly. Consequently, the actin network remained asymmetrically distributed, producing unbalanced steric interactions with microtubules and persistent CS decentering. Robust CS centering was established with ~1000 ERM linkers, where the cortical coverage was sufficient to anchor bulk actin filaments around the cell periphery and prevent their asymmetric redistribution. As a result, microtubules experienced balanced steric resistance from all directions, generating symmetric forces that maintained the CS at the cell center. A further increase to 3500 ERM linkers produced no noticeable improvement, indicating that the cortical anchoring was already saturated at 1000 linkers.

We next examined the effect of ERM turnover by fixing the number of linkers at 1000 and varying the unbinding rate (0.01, 0.1, and 0.3 *s*^−1^), as shown in Fig. 4(h). Increasing the unbinding rate progressively weakened CS centering. At 0.3 *s*^−1^, rapid ERM detachment prevented stable anchoring of bulk actins to the cortex, allowing asymmetric actin redistribution and disrupting the balanced force transmission required for CS centering. In contrast, lower unbinding rates prolonged ERM attachment, stabilized the anchored actin network, and maintained symmetric steric interactions with microtubules. Consequently, CS centering became progressively more stable, with rapid and robust centering observed at an unbinding rate of 0.01 *s*^−1^.

#### 4. Active and passive cross-linkers in bulk actin can not reinforce CS positioning

We next introduced active myosin motors and passive actin cross-linkers into the bulk actin network while maintaining a fixed motor to cross-linker ratio of 1:1. The numbers of motors and cross-linkers were varied in the absence and presence of 3500 ERM linkers, and the resulting CS dynamics are shown in Fig. 4(i), Fig. 4(j), respectively.

As can be seen, the presence and density of ERM linkers remain the primary determinant of CS positioning, whereas variations in the number of myosin motors and passive cross-linkers produce comparatively small changes in CS localization. In the absence of ERM linkers, increasing the density of motors and cross-linkers does not substantially improve the ability of the actin network to maintain the CS near the cell center. This indicates that, although myosin-driven contractility and cross-linker-mediated network organization can modify the internal mechanical properties of the actin network, these effects alone are insufficient to establish a robust positional constraint on the CS.

### C. Influence of centrosomal actin in CS positioning

We next investigated the role of a localized actin meshwork surrounding the centrosome in maintaining its position. Such actin networks have been observed in several cell types, including lymphocytes [14, 38] and dendritic cells [39, 40]. Accordingly, we performed simulations with actin filaments positioned near the centrosome, as shown in Fig. 5(a). Following nucleation, the growing microtubule network exerted pushing forces on the centrosomal actin network, driving it toward the cell periphery. In the absence of ERM-mediated cortical anchoring, the displaced actin network could not establish stable interactions with the cell boundary, and consequently failed to generate restoring forces capable of repositioning the CS toward the geometric center of the cell. As a result, centrosomal actin alone, even at relatively high filament numbers, was insufficient to robustly center the CS Fig. 5(c).

**FIG 5:**
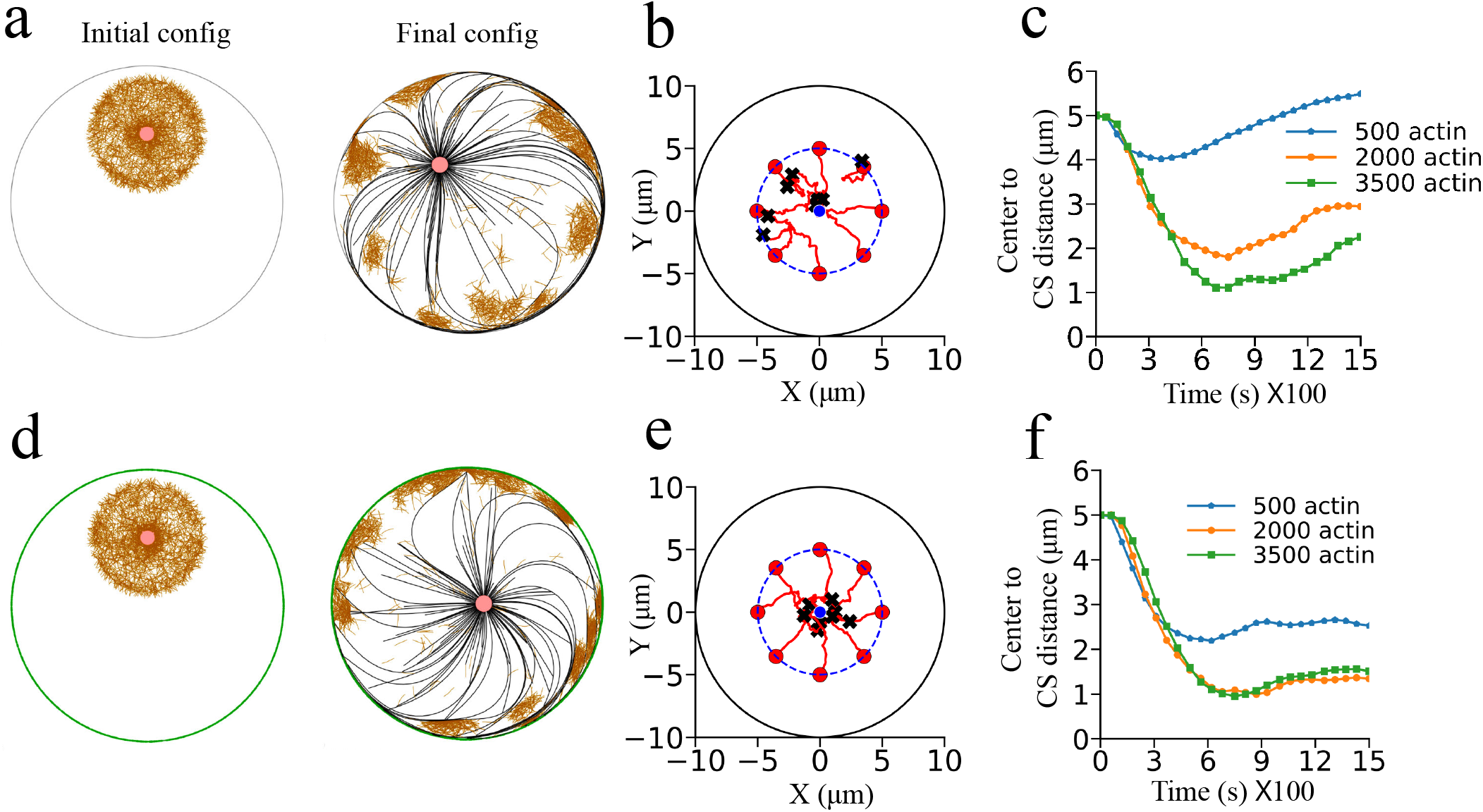
CS positioning with centrosomal actin networks. (a) Initial and final configurations of 2000 actin filaments near the CS without ERM linkers, showing displacement of the network from the cell boundary by growing microtubules. (b) CS trajectories for different initial positions (red circles), showing persistent displacement from the cell center. (c) Mean CS distance from the cell center for different actin filament numbers; increasing actin density promotes centering but does not ensure stable positioning. (d) Initial and final configurations of 2000 actin filaments nucleated near the CS with 3500 ERM linkers (green), showing formation of a mechanically coupled actin cortex. (e) CS trajectories for different initial positions, demonstrating robust centering. (f) Mean CS distance for different actin filament numbers with 3500 ERM linkers. Increasing actin density strengthens and balances cortical force transmission, leading to stable CS positioning near the cell center.

In contrast, when ERM linkers were present at the cortex, the actin filaments reaching the cell boundary became anchored to the membrane, enabling the transmission of forces generated through the interplay between the actin and microtubule networks. Under these conditions, efficient CS centering emerged, indicating that cortical attachment is a critical requirement for this mechanism too, Fig. 5(f).

The efficiency of CS centering was also strongly dependent on the abundance of actin filaments. For actin populations below approximately 2000 filaments, the CS failed to attain a stable central position and remained displaced from the cell center. However, for actin numbers exceeding this threshold, ERM-mediated anchoring together with centrosomal actin consistently promoted CS localization near the cell center. Interestingly, the inclusion of actin cross-linkers and myosin motors did not improve the centering efficiency in the presence of ERM linkers. But when we introduced cross-linkers in the absence of ERM linkers, the off-centering of CS became more pronounced (see Appendix D and E). These findings suggest that robust CS centering in this scenario primarily requires a sufficiently dense centrosomal actin network coupled with effective cortical anchoring.

### D. Positioning of two CSs and microtubule self-organization

We next extended the model to include a second CS. In all cases examined, the two CSs and their associated microtubules spontaneously self-organized into a spindle-like configuration through inter-microtubule steric interactions without the need for actin (Appendix F). This bipolar arrangement resembles the early mitotic spindle and provides a minimal framework for investigating the collective positioning of several CSs.

Using this two-CS model, we examined the role of ERM-mediated membrane-actin coupling in regulating spindle-like organization. Specifically, we investigated how ERM linkers influence CS positioning and spindle organization in the presence of cortical and bulk networks. For each actin architecture, the actin density was varied systematically to determine its effect on CS separation and spindle positioning. This analysis reveals how distinct actin organizations, together with ERM-mediated cortical anchoring, regulate the organization and positioning of spindle-like structures.

#### 1. ERM anchoring facilitates cortical actin regulation of inter-centrosomal distance

We systematically varied the cortical actin numbers (500, 2000, and 3500 filaments) to investigate its effect on the inter-centrosomal (CS-CS) distance. To examine the role of ERM-mediated cortical anchoring, simulations were performed both in the absence of ERM linkers and in the presence of 3500 ERM linkers. A representative comparison for 2000 cortical actin filaments is shown in Fig. 6(a).

**FIG 6:**
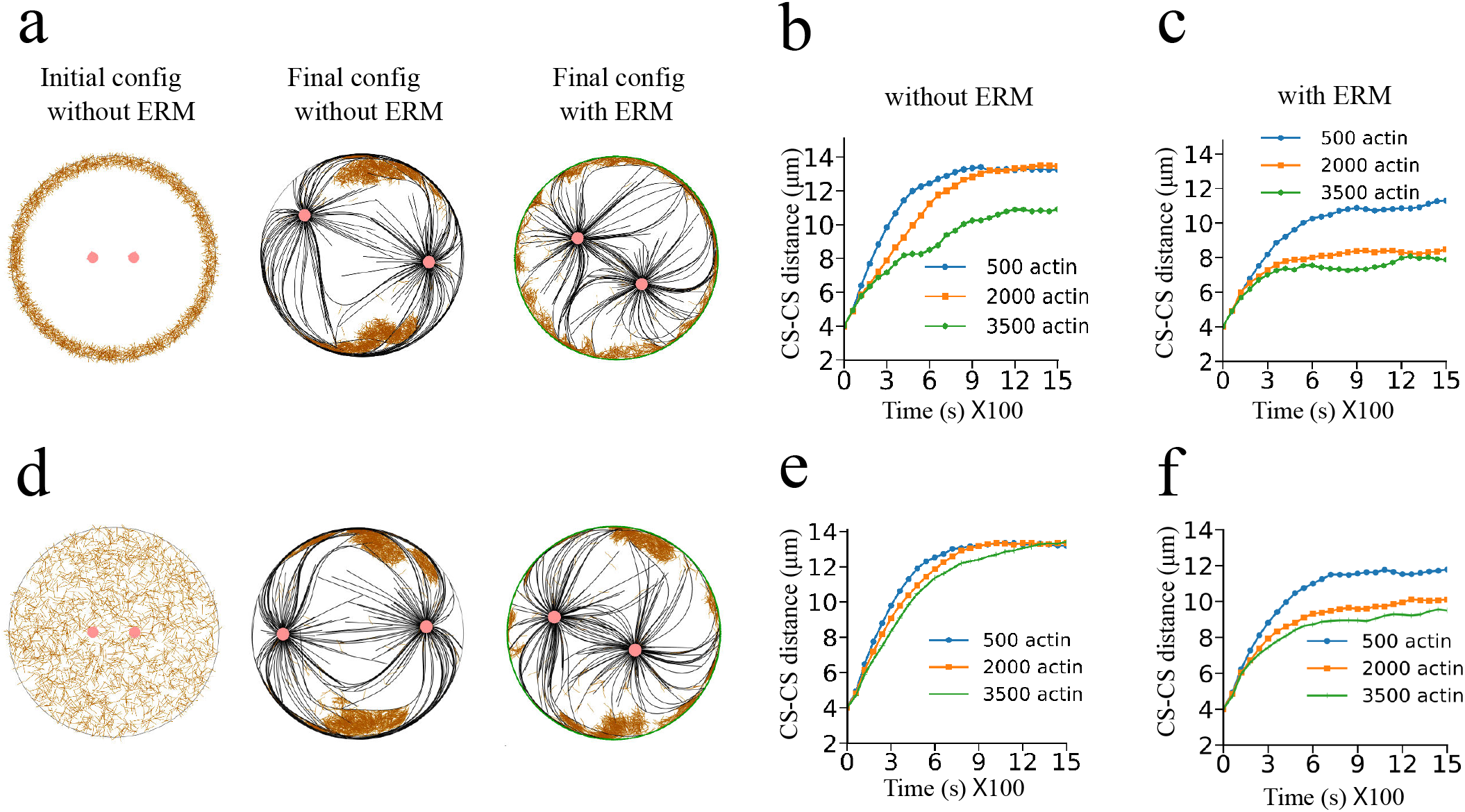
Positioning of two centrosomes and CS–CS distance for different actin configurations. (a–c) Cortical actin networks. (a) Initial and final configurations for 2000 actin filaments without and with ERM linkers. (b) CS–CS distance for different cortical actin numbers without ERM linkers. At low actin densities, growing microtubules sweep actin filaments from their path, allowing the centrosomes to move toward opposite cell poles, while higher actin density increases mechanical resistance and reduces their separation. (c) With ERM linkers, actin–membrane coupling limits their microtubule-driven displacement, with higher actin density further reducing the CS–CS distance. (d–f) Similar results for bulk actin networks. (d) Initial and final configurations for 2000 bulk actin filaments without and with ERM linkers. (e) Without ERM linkers, microtubules can displace a substantial fraction of the bulk actin, allowing larger centrosome separation across the cell. (f) With ERM linkers, even sparse actin–membrane attachment provides resistance to microtubule gliding along the membrane, which increases with actin density and reduces the CS–CS distance.

In the absence of ERM linkers, the CS-CS distance remained nearly unchanged for 500 and 2000 cortical actin filaments, but decreased significantly at 3500 filaments, as summarized in Fig. 6(b). This behavior is consistent with the single-CS results. At low and intermediate actin densities, growing microtubules displaced most cortical actin filaments from their growth paths, resulting in similar mechanical interactions with the cortex and, consequently, comparable CS-CS distances. At a higher actin density of 3500 filaments, however, the cortical actin network could no longer be completely displaced by the growing microtubules. The persistent cortical actin mesh provided stronger steric resistance to microtubule gliding, leading to larger inward pushing forces on the CSs and a reduced equilibrium CS-CS distance.

The corresponding results in the presence of ERM linkers are shown in Fig. 6(c). In contrast to the linker-free case, the CS-CS distance decreased from 500 to 2000 cortical actin filaments. ERM-mediated anchoring prevented cortical actin filaments from being displaced by growing microtubules, allowing the cortical network to remain uniformly distributed along the cell boundary. As the actin density increased, the anchored cortical network generated stronger steric resistance to microtubule gliding, thereby increasing the inward pushing forces on the two CSs and reducing the equilibrium CS-CS distance. A further increase to 3500 actin filaments produced little additional change in the CS-CS distance. With 2000 filaments, the anchored cortical network was already sufficiently dense to maximize the resistance to microtubule gliding, and additional actin filaments contributed only marginally to the force balance.

The analysis shows that ERM-mediated cortical anchoring extends the ability of cortical actin to regulate the inter-centrosomal distance. This regulation is most pronounced at low and intermediate actin densities, whereas at sufficiently high cortical actin density the CS-CS distance becomes largely insensitive to further increases in actin density.

#### 2. Bulk actin regulates inter-centrosomal distance through ERM anchoring

Next, we investigated centrosome positioning in a two-centrosome configuration by considering actin filaments distributed throughout the cytoplasm. We systematically varied the bulk actin numbers (500, 2000, and 3500 filaments) to examine their effects on the inter-centrosomal distance. Representative configurations for 2000 bulk actin filaments are shown in Fig. 6(d).

In the absence of ERM linkers, the CS-CS distance remained nearly unchanged across the range of bulk actin densities, as shown in Fig. 6(e). As microtubules polymerized from the two CSs, they continuously displaced actin filaments from their growth paths through steric interactions. The displaced actin filaments accumulated sparsely within the cell interior rather than being completely pushed to the cell boundary. Consequently, microtubules reached the cortex with little resistance and glided freely along the cell periphery, producing a similar mechanical environment for the CSs irrespective of actin densities considered here. As a result, the equilibrium CS-CS distance remained nearly unchanged.

In contrast, the CS-CS distance became strongly dependent on bulk actin density in the presence of ERM linkers, as shown in Fig. 6(f). Actin filaments that reached the cell boundary through microtubule-driven displacement were captured by ERM linkers and remained anchored to the cortex instead of being redistributed by the growing microtubules. Increasing the bulk or centrosome-associated actin density therefore led to greater cortical accumulation and formation of a denser anchored actin layer. This increased the steric resistance experienced by gliding microtubules and enhanced the inward pushing forces acting on the CSs, leading to a progressive reduction in the equilibrium CS-CS distance. Although a fraction of the actin network remained within the cell interior, the cortically anchored actin layer was sufficient to regulate CS separation.

Thus, we find that increasing bulk actin density alone does not influence the inter-centrosomal distance because the network is continuously reorganized by microtubules. ERM-mediated cortical anchoring fundamentally alters this behavior by retaining actin filaments at the cell boundary, thereby enabling bulk actin density to regulate the equilibrium CS-CS distance.

### E. ERM-mediated actin anchoring tunes spindle length in motor-driven spindles

In the previous section, we showed that a spindle-like structure can form in the double-CS system through steric interactions between microtubules. To obtain a more realistic spindle, we introduced positive end-directed kinesin-5 (Eg5) and negative end-directed motors with similar properties (such as kinesin-14), which generate opposing forces on antiparallel microtubules. Kinesin-5 drives microtubule sliding toward the cell periphery, increasing the CS-CS distance, whereas negative end-directed motors generate inward forces that oppose this separation. Their competition therefore provides a mechanism to establish a finite spindle length.

We fixed the number of negative motors at 500 and varied the number of kinesin-5 motors. The CS-CS distance was found to depend on the motor ratio (Appendix G). We selected a ratio that produced a stable bipolar spindle with a CS-CS distance of approximately 10 *µ*m.

We next introduced cortical and bulk actin networks with filament numbers ranging from 500 to 3500 actin filaments, as shown in Figs. 7(a)-(c) and Figs. 7(d)-(f), respectively. In the presence of ERM linkers, increasing actin density reduced the CS-CS distance in both configurations. Thus, ERM-mediated cortical anchoring continued to regulate spindle length despite the additional forces generated by microtubule motors.

**FIG 7:**
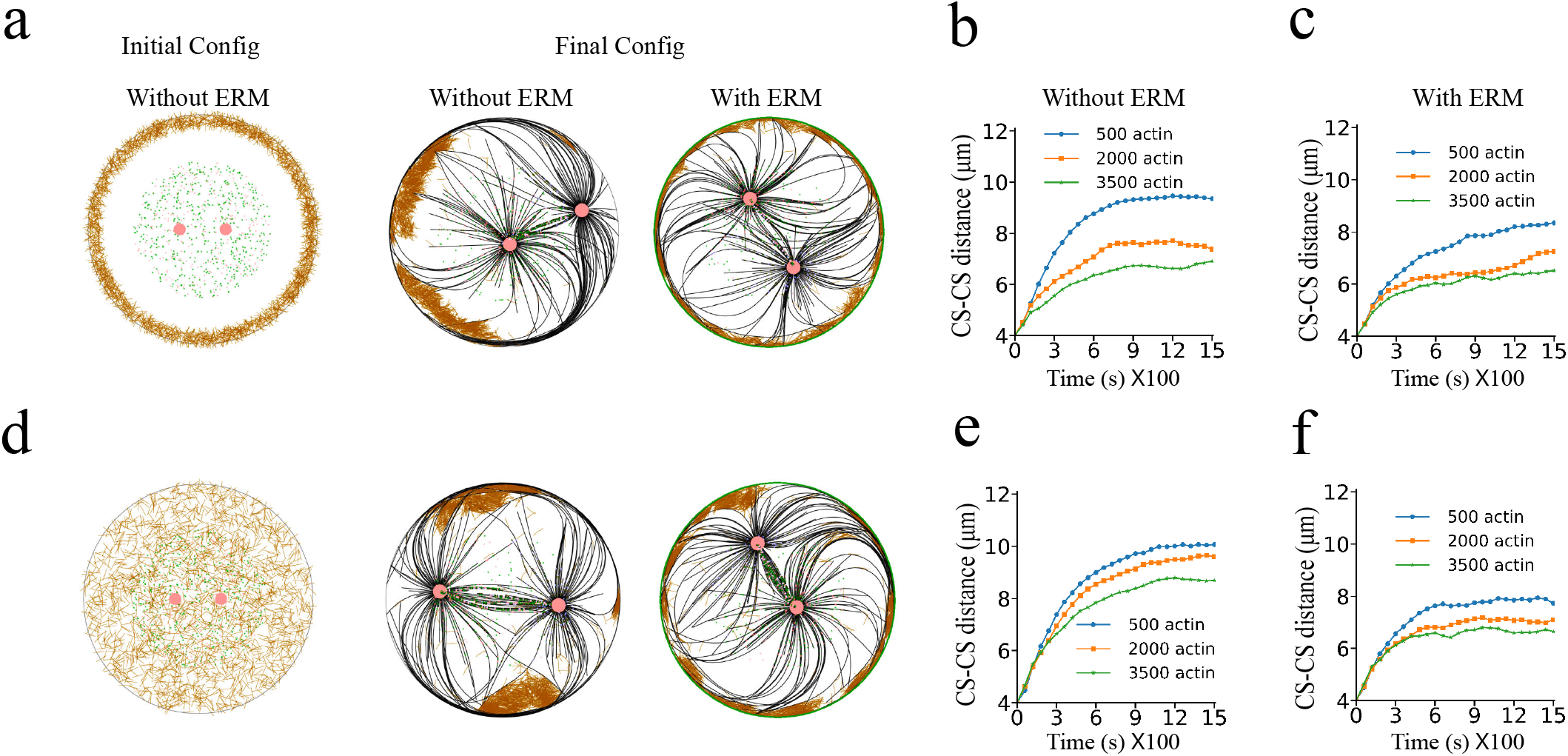
Motor-mediated spindle-like structures with actin networks. (a-c) Cortical actin networks. (a) Initial configuration for 2000 actin filaments without ERM (left) and final configurations without and with ERM linkers. (b) Time evolution of the CS-CS distance for different cortical actin densities without ERM linkers. At low actin density, growing microtubules readily displace the network, whereas increasing the actin number progressively increases resistance and reduces the CS-CS distance. (c) Corresponding results with ERM linkers, where actin-ERM coupling provides additional resistance even at low actin density, with stronger effects at higher actin numbers. (d-f) Bulk actin networks. (d) Initial configuration for 2000 bulk actin filaments without ERM (left) and final configurations without and with ERM linkers. (e) Time evolution of the CS-CS distance without ERM linkers, showing increased resistance and reduced separation with increasing actin density. (f) Corresponding results with ERM linkers, where actin-ERM coupling allows even sparse actin networks to resist microtubule gliding along cell membrane, with further reduction in CS-CS distance at higher actin densities.

The role of the motors was particularly evident in the absence of ERM linkers. For cortical actin, the CS-CS distance decreased from 500 to 2000 actin filaments, as shown in Fig. 7(b). Without motors, steric microtubule interactions rapidly separated the CSs and displaced most cortical actin from the cell boundary. Consequently, 500 and 2000 filaments produced similar CS-CS distances, while only 3500 filaments provided sufficient resistance to reduce spindle length. With kinesin-5 and kinesin-14, the inward force generated by kinesin-14 opposed the outward forces from kinesin-5 and microtubule steric interactions, slowing CS separation. This allowed more cortical actin to remain at the cell boundary, so that 2000 filaments were sufficient to reduce the CS-CS distance. Further increasing the density to 3500 filaments produced little additional change.

A similar effect was observed for bulk actin, as shown in Fig. 7(e). Without motors, rapid CS separation displaced most bulk actin away from the cortex, resulting in nearly identical CS-CS distances across the actin densities. The presence of kinesin-5 and kinesin-14 slowed CS separation and allowed more bulk actin to reach the cell periphery. Increasing the bulk actin density therefore enhanced cortical accumulation and progressively reduced the CS-CS distance, with the strongest effect observed at 3500 filaments.

These results show that antagonistic microtubule motors establish a finite bipolar spindle and alter actin redistribution by slowing CS separation. ERM-mediated anchoring then retains the redistributed actin at the cortex, allowing actin density to further regulate spindle length.

The final CS–CS separation depends on both the actin organization and the presence of ERM linkers and motor proteins. Without motors, the centrosomes remain relatively far apart when ERM proteins are absent, with a separation of about 11–13 *µ*m. Introducing ERM linkers reduces this distance, particularly at higher cortical actin densities. When motor proteins are included, the CS–CS separation becomes smaller overall, and increasing the cortical or bulk actin number leads to a further reduction in the distance. This effect is more pronounced in the presence of ERM linkers, where the separation can reach about 6 *µ*m. A detailed comparison of the CS–CS separation for different actin numbers and combinations of ERM and motor proteins is provided in Appendix H.

## IV. DISCUSSION

In this work, we used numerical results with *Cytosim* [31] to investigate how actin organization and membrane attachment influence interphase CS positioning and mitotic spindle geometry. A schematic of our findings is summarized in Fig. 8. We find that repulsive interactions between growing astral microtubules and cortical actin can promote CS centering [25], but the effectiveness of this mechanism depends on actin density. At low actin density, growing microtubules can displace actin filaments and glide along the cell boundary, producing an asymmetric actin distribution and unbalanced forces on the CS. Increasing actin density strengthens the network and limits microtubule-driven rearrangement, resulting in more balanced forces and improved CS centering.

**FIG 8:**
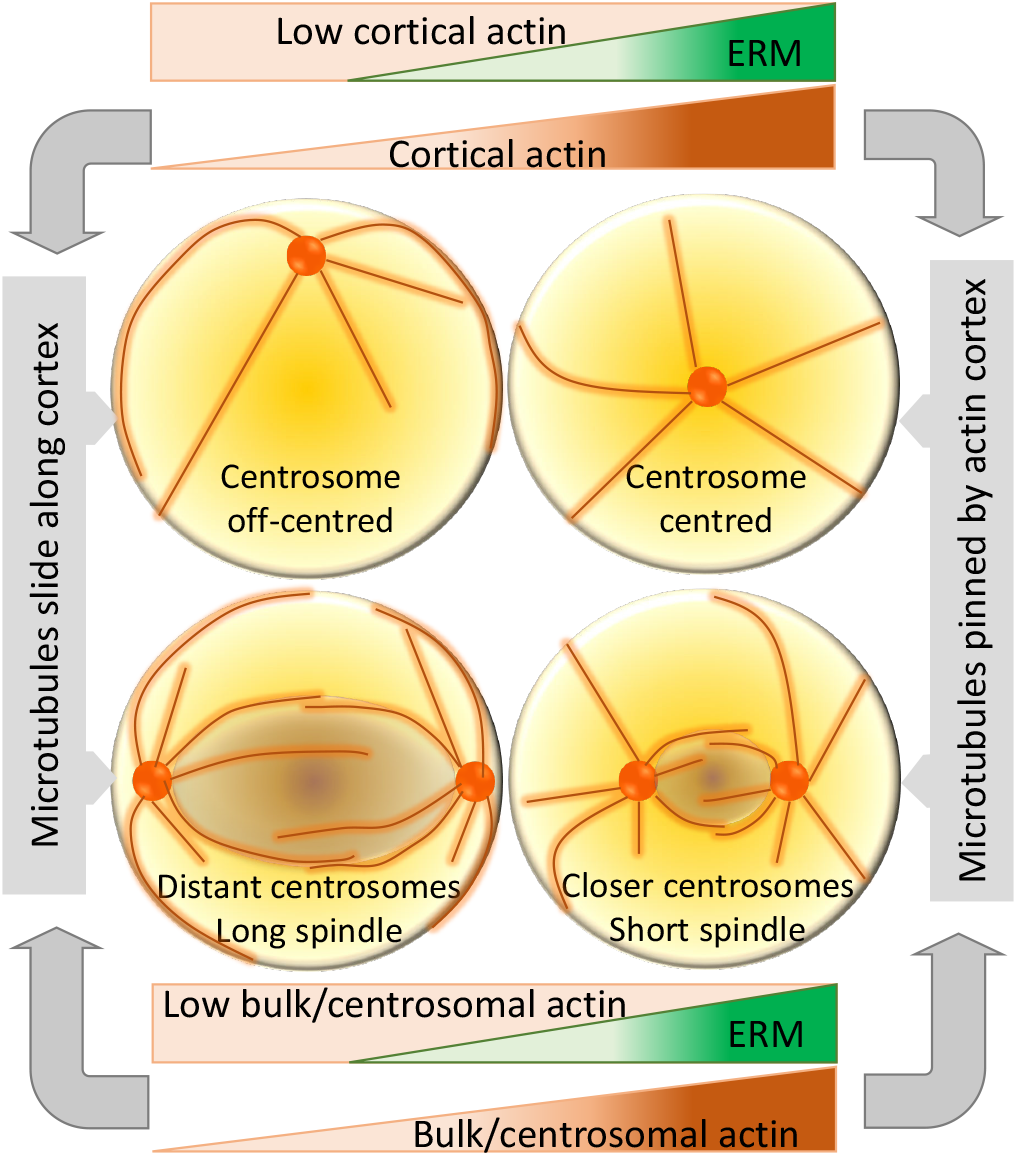
Actin-ERM and microtubule interaction leads to centrosome centering and regulation of spindle length.

**FIG 9:**
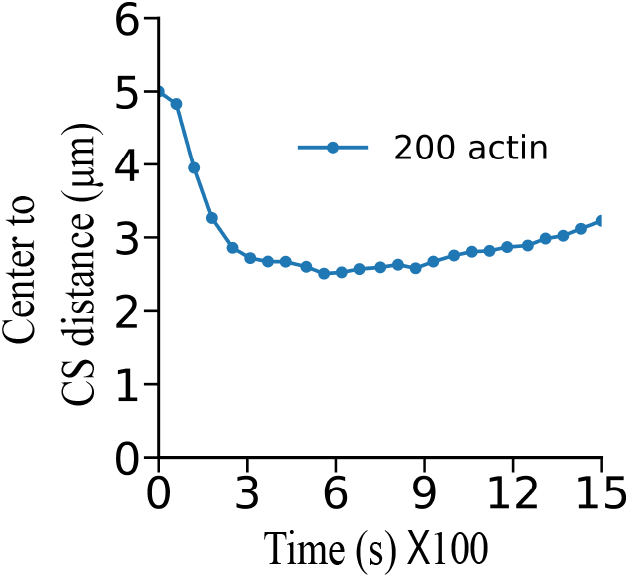
CS positioning in low cortical actin in presence of ERM. At this low cortical actin number (~200), even after being captured by ERM linkers, the actin network is insufficient to generate enough drag on the microtubules to effectively arrest their gliding. Consequently, the forces acting on the CS become asymmetric, resulting in its displacement from the center.

**FIG 10:**
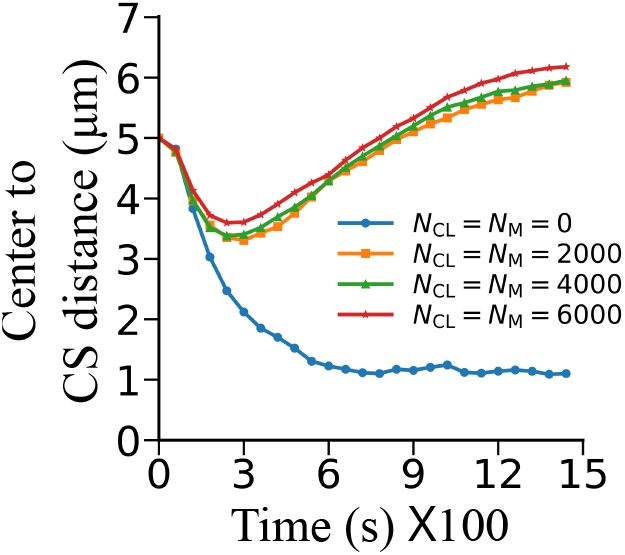
CS positioning for higher cortical actin in presence of cross-linkers and motors without ERM. Without cross-linkers and motors, the relatively high cortical actin density (~2000 filaments) was sufficient to prevent growing microtubules from sweeping the actin filaments away from their paths. This produced a more symmetric force on the CS and allowed it to remain near the cell center. When cross-linkers and myosin motors were introduced, they rearranged the cortical actin network. Since the actin filaments were not strongly anchored to the cell boundary by ERM linkers, the rearranged network could be pushed aside by growing microtubules. This allowed the microtubules to move past the actin network and produced an asymmetric force on the CS, eventually moving it away from the cell center.

**FIG 11:**
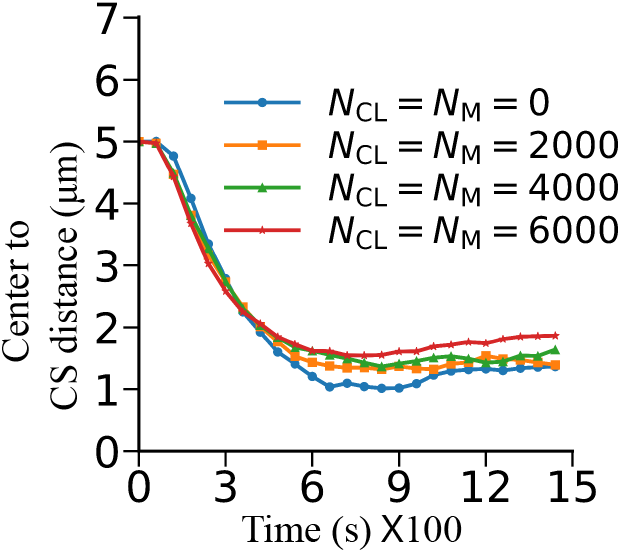
Effect of cross-linkers on CS positioning with ERM for centrosomal actin. ERM proteins stabilize the actin network near the cell boundary, which helps keep the CS close to the cell center. Adding cross-linkers has little effect on the CS positioning at lower numbers. However, at higher cross-linker numbers, the CS moves slightly away from the position observed at zero cross-linkers. This may be due to stronger contraction and reorganization of the actin network, which makes it difficult for the actins to be distributed uniformly along the cell boundary and changes the balance of forces acting on CS.

**FIG 12:**
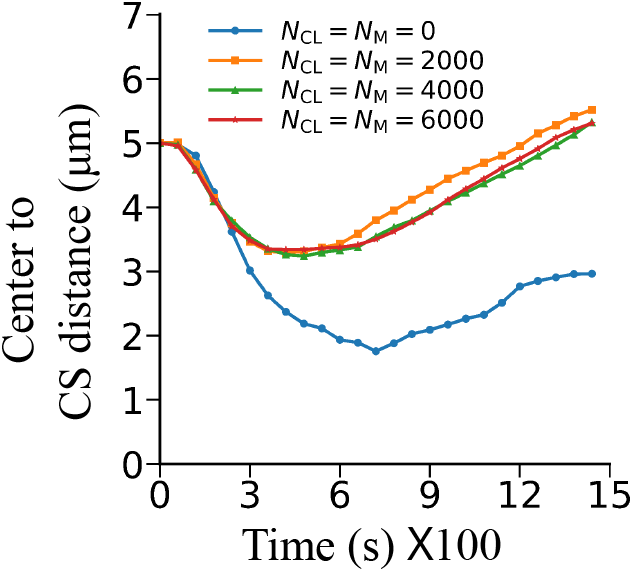
Effect of cross-linkers on CS positioning without ERM for centrosomal actin. Without ERM proteins, CS does not remain close to the cell center. Adding cross-linkers makes off-centering more pronounced. Cross-linkers reorganize and contract the actin network, which leads to a more asymmetric force on CS and further shifts it away from the cell center.

**FIG 13:**
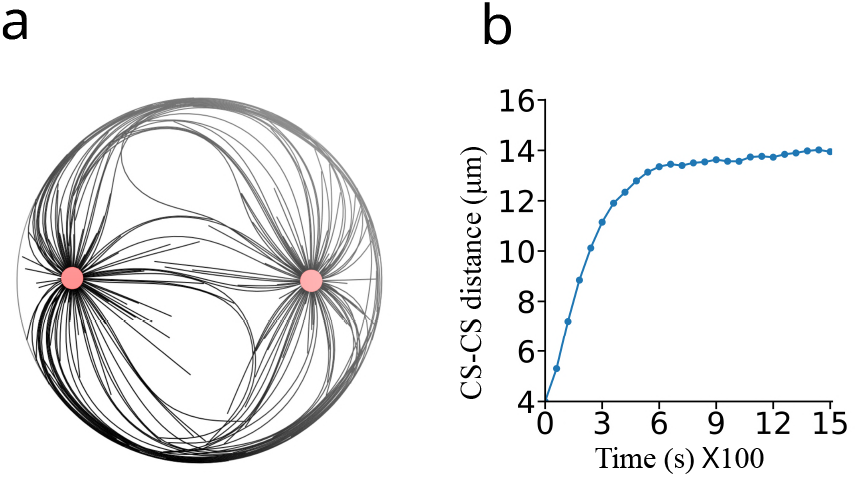
Inter-centrosomal (CS-CS) distance with only microtubule-microtubule (MT-MT) repulsion. (a) Final snapshot of the simulation showing that a spindle-like structure can form through MT-MT repulsion alone. (b) Average CS-CS distance over time. In the absence of actin, microtubules can easily glide along the cell boundary, allowing each centrosome to move toward opposite edges of the cell. At the same time, MT-MT repulsion pushes the two centrosomes apart. As a result, both centrosomes eventually remain near opposite poles, giving a relatively large final CS-CS distance.

**FIG 14:**
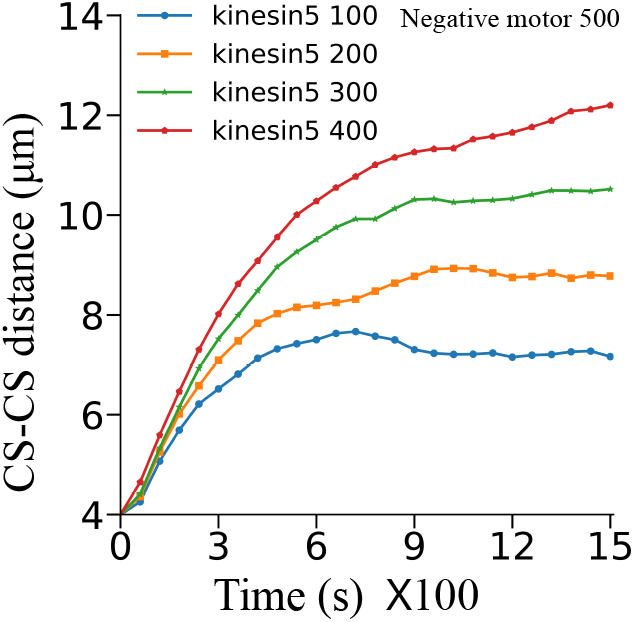
Inter-centrosomal (CS-CS) distance with 500 negative motors and varying numbers of positive motors (kinesin-5). Negative motors attach to two antiparallel microtubules and pull the two centrosomes toward each other, reducing the CS-CS distance, whereas positive motors (kinesin-5) act in the opposite direction and push the centrosomes apart. Keeping the number of negative motors fixed at 500 and increasing the number of positive motors from 100 to 400 progressively increases the CS-CS distance. The results suggest that the balance between the opposing motor activities can regulate the final CS-CS distance.

**FIG 15:**
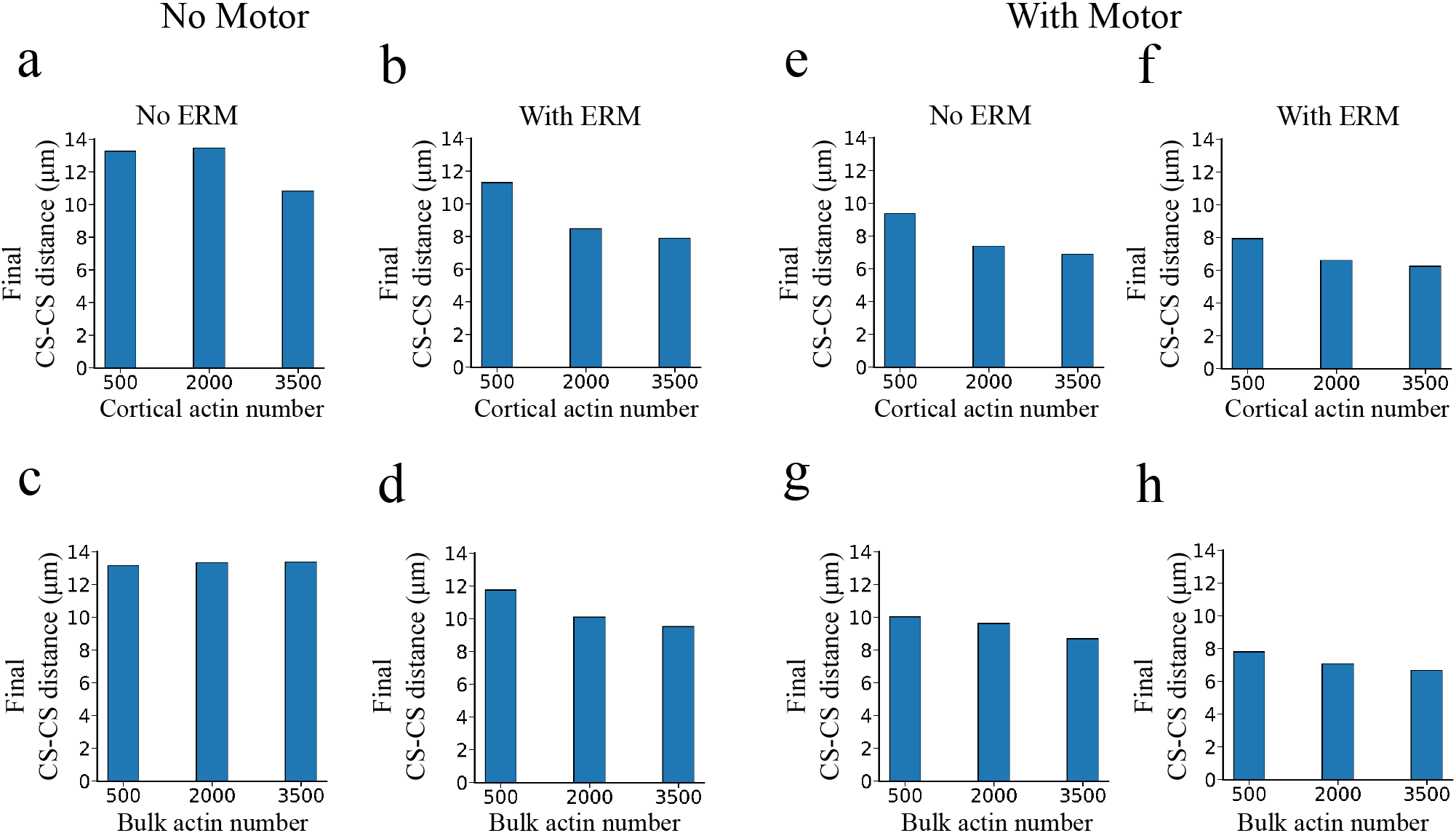
Centrosome–centrosome (CS–CS) distance in the steady state across varying actin numbers in the absence (a–d) and presence (e–h) of motor proteins. Without motors, CS–CS distance remains high (~11–13 *µ*m) when ERM linkers are absent (a, c), while adding ERM linkers noticeably reduces the inter-centrosomal separation, most prominently at higher cortical actin numbers (~8 *µ*m) (b, d). With motor proteins, the overall final CS-CS distance is reduced across all actin numbers (~6–10 *µ*m), with higher cortical (e, f) or bulk (g, h) actin numbers further suppressing separation, an effect enhanced by the presence of ERM (f, h) to reach a minimal distance of ~ 6 *µ*m.

Membrane anchoring through ERM linkers further strengthens this mechanism by coupling the actin network to the cell boundary [20]. This anchoring limits microtubule-driven actin displacement and allows even relatively sparse cortical actin networks to support CS centering. Thus, membrane attachment provides an important mechanical constraint for maintaining cortical organization of actin and stable CS positioning.

We next added passive and active cross-linkers to represent the effects of *α*-actinin and myosin, respectively, simultaneously on the cortical actin network. These proteins can reorganize the network and, in the case of myosin, generate contractile forces [22]. Without ERM linkers, these rearrangements make the initially symmetric network increasingly heterogeneous. Local regions of lower actin density provide less resistance to growing microtubules, allowing them to move along the cortex and displace actin. This can produce unbalanced forces on the CS and perturb stable centering. In contrast, with ERM linkers, the actin network remains mechanically coupled to the cell boundary and is more resistant to microtubule-driven displacement. Consequently, local remodeling by cross-linkers and myosin has a relatively small effect on CS positioning, and centering remains stable.

A similar dependence on membrane anchoring is observed for bulk and centrosome-associated actin networks. Without ERM linkers, growing microtubules can displace these networks even at higher actin densities, leading to asymmetric actin distributions and unbalanced forces on the CS. In the presence of ERM linkers, displaced actin can attach to the cell boundary and provide resistance to microtubule-driven displacement, promoting CS centering. Thus, across the different actin configurations considered here, membrane anchoring has a stronger effect on CS positioning than local actin remodeling by passive or active cross-linkers.

We then extended the model to two centrosomes and found that microtubule interactions alone can generate a bipolar, spindle-like structure. The final CS–CS distance depends on both actin organization and ERM-mediated anchoring. Higher densities of cortical or cortically anchored actin increase resistance to microtubule movement and generate inward forces that limit centrosome separation. Adding opposing motors, such as plus-end-directed kinesin-5 and minus-end-directed kinesin-14, further limits centrosome separation and produces a finite spindle length [41]. Even relatively sparse actin networks can then accumulate near the cell periphery and contribute to spindle-length regulation.

These results provide a mechanical interpretation of recent experimental observations. In particular, the requirement for enough cortical resistance to prevent microtubules from gliding is consistent with in vitro [42], whereas we found membrane-associated actin linkers such as ERM proteins support CS centering at low actin density near the cell cortex. Our simulations suggest that ERM proteins influence CS positioning not only by organizing the cortex but also by setting the mechanical boundary conditions through which microtubule forces are transmitted to the actin network.

More generally, our results show that actin density alone is not sufficient to determine robust CS positioning. Mechanical coupling between the actin network and the cell boundary allows the cortex to resist microtubule forces while the actin network still undergoes local rearrangements. This provides a mechanism through which membrane-bound actin filaments can maintain and stabilize CS position despite changes in the local organization of the actin network during the cell cycle. The two-centrosome simulations further show that the same mechanical interactions can influence spindle geometry. A simple spindle-like structure can form without direct interaction between cortical actin and the spindle, while its geometry can be modulated through microtubule-motor interactions.

Several extensions of this framework would be useful. Incorporating actin turnover would allow us to examine how continuous polymerization and depolymerization affect CS positioning and whether membrane anchoring can maintain centering in a continuously remodeling cortex [43, 44]. It would also be useful to consider force-dependent actin-membrane interactions and spatial variations in actin organization or membrane attachment. These extensions could provide a basis for studying CS positioning under more complex and asymmetric mechanical conditions.

Taken together, we presented a simple mechanical framework for understanding how actin organization and membrane attachment influence CS positioning and spindle geometry. The results emphasize that mechanical coupling to the cell boundary can be as important as actin density itself. Combining these simulations with experimental measurements of actin organization, ERM attachment, and CS dynamics could further clarify how these mechanical interactions contribute to centrosome and spindle organization.

## ACKNOWLEDGMENTS

P.K.B. was supported by a fellowship from the University Grants Commission (UGC), India. A.S. and U.D. acknowledge the Indian Association for the Cultivation of Science, Kolkata, for financial support. R.P. thanks IACS for support and computational facilities.

## Appendix A

## Appendix B: Highly sparse actin network destabilizes centrosome centering in presence of ERM.

## Appendix C: Without ERM, CS-centering is achieved with a very large number of active and passive

## Appendix D: Active and passive cross-linker effect on CS positioning for cs actin case with ERM

## Appendix E: Active and passive cross-linker effect on CS positioning for cs actin case without ERM

## Appendix F: Self-organization of a spindle-like configuration with two centrosomes without actin.

## Appendix G: Spindle length statistics for number of positive and negative motors.

## Appendix H: Comparison of spindle length with respect to ERM, cortical and bulk actin, motor proteins.

**TABLE 1:** Parameters used in the simulations.

| Parameter | Value | Description/reference |
| --- | --- | --- |
| **Microtubules** |  |  |
| Rigidity | $25 \text{ pN}/\mu\text{m}^2$ | Bending rigidity of the microtubule, describing its resistance to bending ([45]) |
| Growth speed | $0.125 \mu\text{m}/\text{s}$ | Rate at which a growing microtubule elongates ([8]) |
| Shrinkage speed | $0.266 \mu\text{m}/\text{s}$ | Rate at which a shrinking microtubule shortens ([8]) |
| Rescue rate | $0.067 \text{ s}^{-1}$ | Rate of transition from microtubule shrinkage to growth ([8]) |
| Catastrophe rate | $0.03, 0.04 \text{ s}^{-1}$ | Rates of transition from microtubule growth to shrinkage for unloaded and stalled states |
| **Centrosome** |  |  |
| Radius | $0.55 \mu\text{m}$ | Radius of the centrosome |
| Effective viscosity | $200 \text{ pN s}/\mu\text{m}^2$ | To match the mobility of centrosome as measured in experiments ([11]) |
| First anchoring stiffness | $500 \text{ pN}/\mu\text{m}$ | Stiffness of first link between MT minus ends and centrosome center ([12]) |
| Second anchoring stiffness | $500 \text{ pN}/\mu\text{m}$ | Stiffness of second link between MT minus ends and a point on the centrosome periphery; the non-zero high value prevents the MTs from pivoting ([12]) |
| Number of microtubules | 100 | Adopted from ([11]) |
| **Actin** |  |  |
| Rigidity | $0.05 \text{ pN}/\mu\text{m}^2$ | Adopted from ([46]) |
| Length | $1.0 \mu\text{m}$ | We used actins of fixed length |
| **Motor** |  |  |
| Binding rate | $5 \text{ s}^{-1}$ | Rate at which myosin heads bind to nearby actin filaments ([47]) |
| Binding range | $50 \text{ nm}$ | Adopted from ([48]) |
| Unbinding force | $3.65 \text{ pN}$ | Force required to detach from attached filaments |
| Max. speed | $0.3 \mu\text{m s}^{-1}$ | Speed of the motor head towards the positive end ([49]) |
| Stall force | $2 \text{ pN}$ | Force at which the motor stops moving under load |
| Stiffness | $100 \text{ pN}/\mu\text{m}$ | Spring constant of the link between two motor heads ([48]) |
| Length | $30 \text{ nm}$ | Rest length of the link ([47]) |
| **Crosslinker** |  |  |
| Stiffness | $100 \text{ pN}/\mu\text{m}$ | Spring constant of the link between two crosslinker heads, same value used as for motors |
| Length | $40 \text{ nm}$ | Rest length of the link, α-actinin length ([50]) |
| **ERM** |  |  |
| Stiffness | $1000 \text{ pN}/\mu\text{m}$ | The ERM–actin spring constant was set to $1000 \text{ pN}/\mu\text{m}$ for a stiff bond. Using $100 \text{ pN}/\mu\text{m}$ gave similar results. |
| **Confinement space** |  |  |
| Cell radius | $10 \mu\text{m}$ | Radius of the circular cell boundary |
| Confinement stiffness | $500 \text{ pN}/\mu\text{m}$ | Confinement of filaments and other elements inside the cell ([12]) |

## References

[1] M. Bornens, Open Biology 8, 180052 (2018).

[2] A. Desai and T. J. Mitchison, Annual review of cell and developmental biology 13, 83 (1997).

[3] I. Stötzel, A.-K. Weier, A. Sarkar, S. Som, L. Bach, P. Konopka, E. Miková, S. Ghosh, J. Böthling, M. Hom-rich, et al., Nature Communications 17, 536 (2026).

[4] A. Sarkar, H. Rieger, and R. Paul, Biophysical journal116, 2079 (2019).

[5] A. Sarkar, A. Jana, A. Agashe, J. Wang, R. Kapania,N. S. Gov, J. G. DeLuca, R. Paul, and A. S. Nain, PNAS nexus 4, pgaf201 (2025).

[6] J. L. Badano, T. M. Teslovich, and N. Katsanis, Nature Reviews Genetics 6, 194 (2005).

[7] A. Jana, A. Sarkar, H. Zhang, A. Agashe, J. Wang,R. Paul, N. S. Gov, J. G. DeLuca, and A. S. Nain, Proceedings of the National Academy of Sciences 120, e2120536120 (2023).

[8] A. Burakov, E. Nadezhdina, B. Slepchenko, and V. Rodionov, The Journal of cell biology 162, 963 (2003).

[9] S. Som, S. Chatterjee, and R. Paul, Physical Review E 99, 012409 (2019).

[10] A. Mallick, A. Sarkar, and R. Paul, Indian Journal of Physics 96, 2667 (2022).

[11] J. Zhu, A. Burakov, V. Rodionov, and A. Mogilner, Molecular biology of the cell 21, 4418 (2010).

[12] G. Letort, F. Nedelec, L. Blanchoin, and M. Théry, Molecular biology of the cell 27, 2833 (2016).

[13] U. Euteneuer and M. Schliwa, Journal of Cell Biology101, 96 (1985).

[14] F. Farina, J. Gaillard, C. Guérin, Y. Couté, J. Sillibourne,L. Blanchoin, and M. Théry, Nature cell biology 18, 65 (2016).

[15] T. M. Svitkina, Trends in cell biology 30, 556 (2020).

[16] D. Obino, F. Farina, O. Malbec, P. J. Sáez, M. Maurin,J. Gaillard, F. Dingli, D. Loew, A. Gautreau, M.-I. Yuseff,et al., Nature communications 7, 10969 (2016).

[17] S. Yamamoto, J. Gaillard, B. Vianay, C. Guerin,M. Orhant-Prioux, L. Blanchoin, and M. Théry, The EMBO Journal 41, EMBJ2022111631 (2022).

[18] S. L. Prosser and L. Pelletier, Nature Reviews Molecular Cell Biology 18, 187 (2017).

[19] R. G. Fehon, A. I. McClatchey, and A. Bretscher, Nature Reviews Molecular Cell Biology 11, 276 (2010).

[20] S. Tsukita, S. Yonemura, and S. Tsukita, Trends in biochemical sciences 22, 53 (1997).

[21] A. Bretscher, K. Edwards, and R. G. Fehon, Nature reviews Molecular cell biology 3, 586 (2002).

[22] M. Murrell, P. W. Oakes, M. Lenz, and M. L. Gardel, Nature Reviews Molecular Cell Biology 16, 486 (2015).

[23] H. Ennomani, G. Letort, C. Guerin, J.-L. Martiel, W. Cao,F. Nedelec, E. M. De La Cruz, M. Thery, and L. Blanchoin, Current Opinion in Cell Biology 50, 79 (2018).

[24] M. R. Hannaford and N. M. Rusan, Journal of Cell Biology223, e202311140 (2024).

[25] A. V. Burakov and E. S. Nadezhdina, Cells 9, 1351 (2020).

[26] T. E. Holy, M. Dogterom, B. Yurke, and S. Leibler, Proceedings of the National Academy of Sciences 94, 6228 (1997).

[27] H.-Y. Wu, G. Kabacaoğlu, E. Nazockdast, H.-C. Chang, M. J. Shelley, and D. J. Needleman, Nature Physics 20, 157 (2024).

[28] L. Laan, N. Pavin, J. Husson, G. Romet-Lemonne,M. Van Duijn, M. P. López, R. D. Vale, F. Jülicher,S. L. Reck-Peterson, and M. Dogterom, Cell 148, 502 (2012).

[29] K. Kimura and A. Kimura, Bioarchitecture 1, 74 (2011).

[30] A. Schaeffer, S. Buracco, M. Gazzola, M. Gelin, B. Vianay,C. de Pascalis, L. Blanchoin, and M. Théry, Journal of Cell Biology 224, e202405126 (2025).

[31] F. Nedelec and D. Foethke, New Journal of Physics 9, 427 (2007).

[32] T. Mitchison and M. Kirschner, nature 312, 237 (1984).

[33] T. P. Kole, Y. Tseng, I. Jiang, J. L. Katz, and D. Wirtz, Molecular biology of the cell 16, 328 (2005).

[34] O. Esue, Y. Tseng, and D. Wirtz, PLoS One 4, e4411 (2009).

[35] Y. Bashirzadeh, H. Moghimianavval, and A. P. Liu, Iscience 25 (2022).

[36] M. Murrell, P. W. Oakes, M. Lenz, and M. L. Gardel, Nature reviews Molecular cell biology 16, 486 (2015).

[37] S. Wang and P. G. Wolynes, Proceedings of the National Academy of Sciences 109, 6446 (2012).

[38] D. Inoue, D. Obino, J. Pineau, F. Farina, J. Gaillard,C. Guerin, L. Blanchoin, A.-M. Lennon-Duménil, and M. Théry, The EMBO journal 38, EMBJ201899630 (2019).

[39] M. T. Schmitt, J. Kroll, M. J. Ruiz-Fernandez,R. Hauschild, S. Ghosh, P. Kameritsch, J. Merrin, J. Schmid, K. Stefanowski, A. W. Thomae, et al., Science advances 11, eadx4047 (2025).

[40] A.-K. Weier, M. Homrich, S. Ebbinghaus, P. Juda, E. Miková, R. Hauschild, L. Zhang, T. Quast, E. Mass,A. Schlitzer, et al., Journal of Cell Biology 221, e202107134 (2022).

[41] C. Hentrich and T. Surrey, Journal of Cell Biology 189, 465 (2010).

[42] A. J. Jimenez, A. Schaeffer, C. De Pascalis, G. Letort, B. Vianay, M. Bornens, M. Piel, L. Blanchoin, and M. Théry, Current Biology 31, 1206 (2021).

[43] B. L. Goode, J. Eskin, and S. Shekhar, Journal of Cell Biology 222, e202309021 (2023).

[44] M.-F. Carlier and S. Shekhar, Nature Reviews Molecular Cell Biology 18, 389 (2017).

[45] F. Gittes, B. Mickey, J. Nettleton, and J. Howard, The Journal of cell biology 120, 923 (1993).

[46] W. Luo, C.-h. Yu, Z. Z. Lieu, J. Allard, A. Mogilner, M. P. Sheetz, and A. D. Bershadsky, Journal of Cell Biology 202, 1057 (2013).

[47] J. P. Robblee, A. O. Olivares, and M. Enrique, Journal of Biological Chemistry 279, 38608 (2004).

[48] H. Ennomani, G. Letort, C. Guérin, J.-L. Martiel, W. Cao,F. Nedelec, E. M. De La Cruz, M. Thery, and L. Blanchoin, Current Biology 26, 616 (2016).

[49] R. S. Rock, S. E. Rice, A. L. Wells, T. J. Purcell, J. A. Spudich, and H. L. Sweeney, Proceedings of the National Academy of Sciences 98, 13655 (2001).

[50] D. H. Wachsstock, W. Schwartz, and T. D. Pollard, Biophysical journal 65, 205 (1993).

